# Real-time computation of the TMS-induced electric field in a realistic head model

**DOI:** 10.1101/547315

**Authors:** Matti Stenroos, Lari M. Koponen

## Abstract

Transcranial magnetic stimulation (TMS) is often targeted using a model of TMS-induced electric field (E). In such navigated TMS, the E-field models have been based on spherical approximation of the head. Such models omit the effects of cerebrospinal fluid (CSF) and gyral folding, leading to potentially large errors in the computed E-field. So far, realistic models have been too slow for interactive TMS navigation. We present computational methods that enable real-time solving of the E-field in a realistic five-compartment (5-C) head model that contains isotropic white matter, gray matter, CSF, skull and scalp.

Using reciprocity and Geselowitz integral equation, we separate the computations to coil-dependent and -independent parts. For the Geselowitz integrals, we present a fast numerical quadrature. Further, we present a moment-matching approach for optimizing dipole-based coil models.

We verified and benchmarked the new methods using simulations with over 100 coil locations. The new quadrature introduced a relative error (RE) of 0.3–0.6%. For a coil model with 42 dipoles, the total RE of the quadrature and coil model was 0.44–0.72%. Taking also other model errors into account, the contribution of the new approximations to the RE was 0.1%. For comparison, the RE due to omitting the separation of white and gray matter was >11%, and the RE due to omitting also the CSF was >23%.

After the coil-independent part of the model has been built, E-fields can be computed very quickly: Using a standard PC and basic GPU, our solver computed the full E-field in a 5-C model in 9000 points on the cortex in 27 coil positions per second (cps). When the separation of white and gray matter was omitted, the speed was 43–65 cps. Solving only one component of the E-field tripled the speed.

The presented methods enable real-time solving of the TMS-induced E-field in a realistic head model that contains the CSF and gyral folding. The new methodology allows more accurate targeting and precise adjustment of stimulation intensity during experimental or clinical TMS mapping.

## 1. Introduction

In transcranial magnetic stimulation (TMS) [1], a brief current pulse in a coil gives rise to a time-varying magnetic field that causes an electric field (E-field), and the E-field stimulates the brain. The E-field consists of two parts: the primary E-field 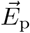 induced by the changing magnetic field causes a current distribution in the head, and this current is affected by changes of electric conductivity, leading to charge buildup at conductivity interfaces. These charges give rise to a secondary electric field 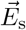, the total E-field thus being 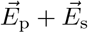.

In navigated TMS, the stimulation is targeted and its intensity is set using a model of TMS-induced E-field. For this, we need a volume conductor model (VCM) that characterizes the conductivity distribution of the head. While simulation and offline TMS studies use realistically-shaped highly detailed VCMs [2, 3, 4, 5], in experimental navigated TMS the head has so far been modeled as a spherically symmetric conductor that is fitted to approximate either the shape of the skull under the coil or the whole skull. Spherical models omit the effects of well-conducting cerebrospinal fluid (CSF) [2, 6] and may fail in approximating the effects of skull at regions of changing skull curvature [7].

To optimally benefit from the E-field model in navigation, the E-field should be computed in near real time, so that the operator can see the estimated field pattern while moving the TMS coil. With finite-element methods used for solving realistic VCMs in, for example, [4, 5] and recently proposed approaches that apply finite differences [8] and boundary elements [9, 10], computation of the E-field takes of the order of tens of seconds to minutes for one coil position.

In this paper, we present and benchmark computational methods that allow fast solving the E-field in a realistic VCM that contains realistic CSF and gray matter and isotropic white matter, tens of coil positions per second. Our approach is based on the reciprocity relation [11, 12] and surface-integral solution of the magnetic field in a VCM [13]. The same idea has earlier been applied in, for example, [7]. Here, we present a formulation optimized for repeated computations, and two novel techniques to speed up the computations: a simple-but-adequate numerical quadrature for surface integrals and a moment-matching approach to optimize the coil model for rapid computations. We verify and benchmark the new methods using simulations in four-compartment and five-compartment models, showing that together our new methods enable the real-time computation of E-field in a highly realistic VCM. The fast quadrature is applicable to any surface-based approach in isotropic VCMs, and the coil-modeling technique can be used in any computational environment that models coils using a set of dipoles.

## 2. Theory

### 2.1. Reciprocity relation

Let there be a system of conductors. Let primary current distribution 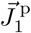 give rise to electric field 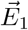. Correspondingly,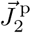 gives rise to 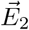. Then, assuming the head electrically resistive and small compared to the wavelength of electromagnetic waves, we get the reciprocity relation [11]

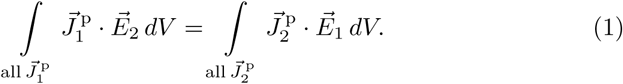

Letting 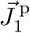 be a dipolar current element 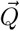 at 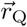, i.e. 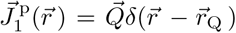, and 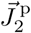 a current loop (coil) with current *I* flowing in a thin wire *c*, we get [12]

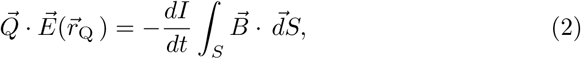

where 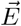 is the electric field generated by *dI=dt*, 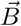 is the magnetic field produced by 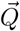, and *c* = *∂S* is the boundary of *S*. Setting an unit-current dipole 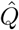 to 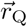 and computing its magnetic flux through the coil, we get the corresponding component of 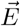 in 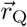, as illustrated in Fig. 1.

**Figure 1:**
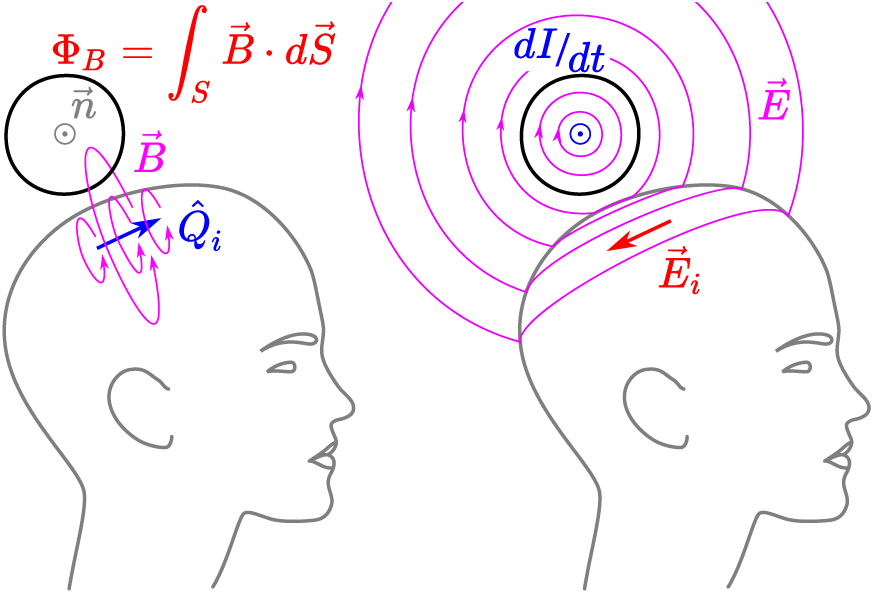
The reciprocity relation of Eq. 2: Computing the magnetic flux Φ_B_, where 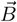 is the magnetic field due to a unit-current dipole 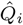, we obtain the relationship between the rate of change in coil current (dI/dt) and one component of the induced E-field 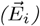 at the location of the current dipole. Note that 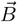 is only evaluated inside the coil, and that 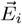 is obtained without evaluating the E-field 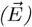 elsewhere.

### 2.2. Forward problem for primary current distribution in the head

When computing the TMS-induced E-field reciprocally, we solve the traditional biomagnetic forward problem: what is the 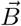 outside the head, when there is a primary current distribution 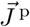 inside the head? Here we present the basic equations for solving this problem using surface integral approach and discretized boundary potentials and collect the solution to an efficient form.

Consider a primary current distribution in a piece-wise homogeneous isotropic volume conductor that has *K* boundary surfaces separating the compartments of different conductivities. Let superscripts in italics label boundary surfaces. The magnetic field in or outside of the conductor is [13]

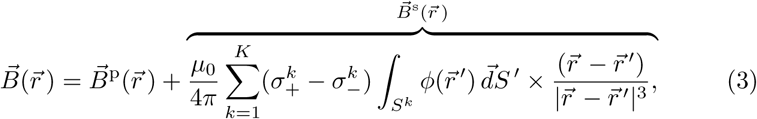

where 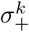 and 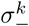 are the conductivities outside and inside of boundary *k*, 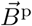 is the primary magnetic field due to 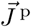, *ϕ* is the electric potential generated by the charge density associated with the 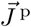, and 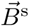 is the secondary magnetic field due to conductivity inhomogeneities in the head.

Now discretize the conductivity boundaries into a set of vertices and triangular elements; boundary *k* has *N* ^*k*^ vertices, and the total number of vertices is 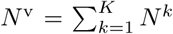. Express the boundary potentials *ϕ*^*k*^ using basis functions: 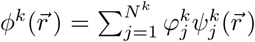, where 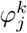 is the potential in the vertex *j* of boundary *k* (target vertex), and 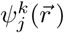 is a linear basis function that is spanned on the member triangles of the target vertex and gets value of 1 in the target vertex and value of 0 in the other vertices^1^. We get

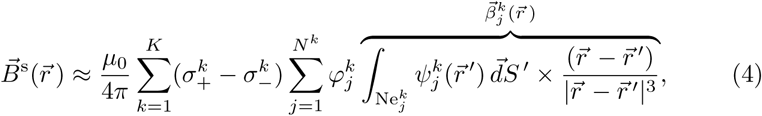

where 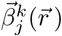 is the integral that needs to be computed repeatedly for all elements and field points, when the TMS coil is moved. When the boundary potentials are solved numerically using the (boundary) basis functions 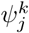, Eq. 3 does not introduce any numerical error, as no interpolation or additional discretization is needed.

To compute the magnetic flux through a coil, we need to set up a numerical integration scheme. For this purpose, we compute the normal component 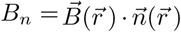 in *N*^*c*^ quadrature points in the coil. For any source distribution, we then get the matrix form

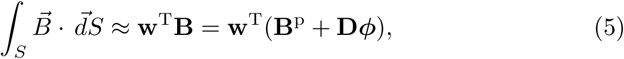

where **w**^T^ (1×*N* ^c^) contains coil quadrature weights, **B**^p^ (*N* ^c^ ×1) contains 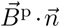 in the quadrature points, **D** (*N* ^c^ × *N* ^v^) contains 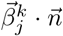 for the coil quadrature points and all basis functions, and ***ϕ*** (*N* ^v^× 1) contains 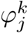 in all model vertices and all the remaining constants and material parameters of Eq. 4.

To compute TMS-induced 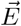 using Eq. 2, we need to solve Eq. 5 for dipole sources placed all over the cortical region of interest (ROI, with *N* ^ROI^ dipoles). The accuracy and computational cost of the solution depend on:

1. the accuracy of the volume conductor model and ***ϕ***; the discretization and number of vertices *N* ^v^,
2. the method used for computing 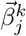,
3. the accuracy and number of quadrature points *N* ^c^ of the coil model, and
4. the extent and discretization density *N* ^ROI^ of the ROI.

Potential ***ϕ*** does not depend on the TMS coil. It can thus be computed in advance for the region of interest, using the desired level of detail and numerical approach.

### 2.3. Fast quadrature for computing secondary magnetic field

When accurately computing the secondary magnetic field 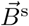 using Eq. 5, we need to evaluate *N* ^v^ surface integrals, each spanned in the neighborhood of one vertex. Typically the computation is carried out triangle-wise, using linear shape functions that have value of 1 in one vertex and value of 0 in the other vertices of the triangle. For these shape functions, we have a closed-form analytical solution of 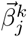 [14]. In applications, where no real-time modeling is needed (for example, magnetoencephalography and magnetocardiography), that approach works excellently. For the real-time TMS approach it might, however, become a bottleneck, so we will apply numerical integration.

In numerical integration over a meshed surface, each element is typically represented by a set of quadrature points 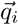 and weights *w*_*i*_: for one element,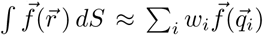. With flat triangular elements, Gaussian quadrature with either 7 or 13 points per triangle is a typical choice. As each vertex has on the average six member triangles, a 7-point quadrature would result in evaluating the integrand of 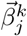 in approximately 42 points. Each triangle belonging to three vertices, the approximate number of quadrature points per vertex would then be 14. We will, however, take one step further and represent 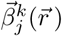 using only one quadrature point per vertex neighborhood.

Let vertex 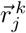 have *N* member triangles *T*_*i*_ that have normal vectors 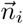 and areas *A*_*i*_. The basis function *ψ* is then expressed using *N* shape functions *s*_*i*_. For each 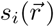, find its center of mass 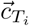 and compute the dipole moment 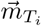 of 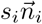. Then, to get the equivalent dipole for the whole 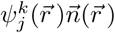, compute the weighted center of mass 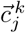 and the total dipole moment 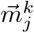:

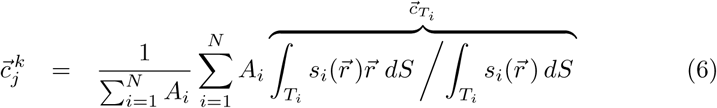

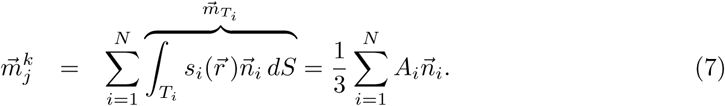

Thus

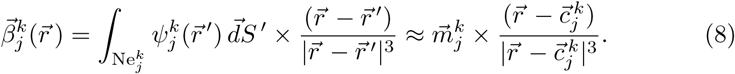

This quadrature is considerably simpler and lighter to compute that the closed-form solution that contains trigonometry, logarithms, and several times more elementary vector operations.

### 2.4. Coil model

To obtain the E-field, we numerically compute the integral on the right hand side of Eq. 2. The coil is represented as a set of (oriented) quadrature points as in the previous section. Or, thinking the other way, the current loops of the coil are represented as a set of magnetic dipoles (see, e.g., [15]). A set of dipoles that represents a TMS coil can be generated by discretizing the volume where the coil windings lie into smaller sub-volumes [15, 16, 7]; similar approaches are also used with permanent magnets [17]. As the TMS coil is relatively large and close to the head, this approach, however, requires a large number of dipoles for accurate representation of the windings. For example, in [15], the Magstim 70mm Double Coil (P/N 9925) whose windings span a volume of 176 × 88 × 7 mm^3^ was represented with 2712 dipoles in three identical layers. For real-time computation, we would prefer a much lower number of dipoles—of the order of at most one hundred.

For a closed current loop *c* with current *I*, the magnetic dipole moment is 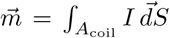, where *A* is any open surface bounded by *c* [18]. For a unit-current coil whose (zero-height) windings lie on such a surface, the magnetic dipole moment density *f*_0_ at each point (*x*_1_, *x*_2_) on *A*_coil_ thus equals the number of turns surrounding that point. With non-zero height of the windings, the dipole moment density becomes *f* (*x*_1_, *x*_2_, *x*_3_) = *f*_0_(*x*_1_, *x*_2_)*g*(*x*_1_, *x*_2_, *x*_3_), where *g* characterizes the spread of current along the height of the windings. This kind of density functions can be characterized by geometric moments [19].

Now consider a planar coil in *xy* plane, with current spread evenly across the height of the windings *h*. The dipole moment density is *f* = *f*_0_(*x, y*)*/h*, and all dipoles point at 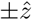 direction. The geometric moment *µ* of the order (*n*_x_, *n*_y_, *n*_z_) is

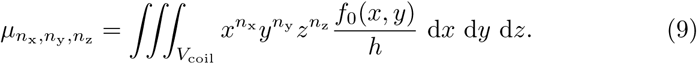

As *f* = *f*_0_*/h* does not depend on *z*, the calculation can be separated into two parts,

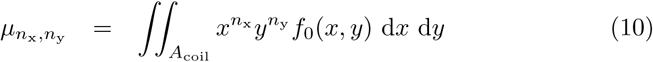

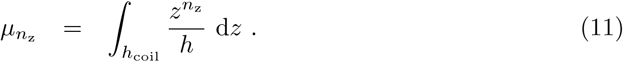

Consider first the simpler *z* direction. Place the coil symmetrically around the xy plane. From Eq. 11 we see that all odd moments are zero; for even *n*_z_, we get

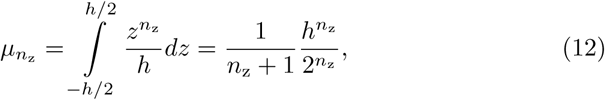

while for a discrete set of dipoles in locations *z*_*i*_ we get 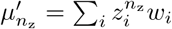, where *w*_*i*_ is the weight of each dipole. For odd moments 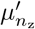 to vanish, we place the dipoles symmetrically around the xy plane in locations ±*z*_*i*_ (*i* > 0) and allow a dipole on xy plane, *z*_0_ = 0. We then have

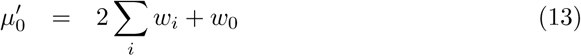

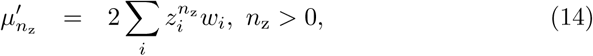

where *w*_0_ = 0 for even number of dipoles. With *I* dipoles, we thus have *I* free parameters, which we can solve analytically from a system of equations

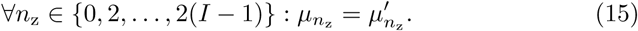

For the *x* and *y* directions, the (*n*_x_, *n*_y_)-order moment of a discrete set of dipoles is

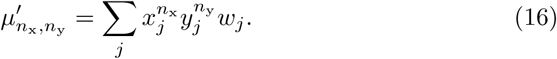

Coils currently used for navigated TMS are either circular or figure-of-eight type. A figure-of-eight coil is commonly expressed as a set of circular loops. Consider thus a circular loop, radius *r*, at origin. The geometrical moments for such a loop can be computed in closed form with, e.g., Mathematica software (version 10.3.0, www.wolfram.com). When both *n*_x_ and *n*_y_ are even, the (*n*_x_, *n*_y_)-order moment for the corresponding *f* (*x, y*) is

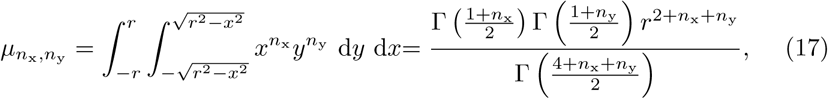

where Γ is the gamma function. If either *n*_x_ or *n*_y_ is odd, 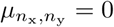.

To reproduce the moments of a coil with dipoles, we place *n* dipoles per ring in 0–2 rings so that there is symmetry along at least one axis and *n*-fold rotational symmetry. In addition, we allow a point at the origin—see Figure 2 for an example placement. In Eq. 17, the ratios between (*n*_x_, *n*_y_)-order moments of a given degree, *N* = *n*_x_ + *n*_y_, are rational numbers. For sufficient *n*, the ring-like structure will, due to symmetry, reproduce these ratios for all degrees up to *N*. Thus, it will always match either none or all moments of degree *N* or below, which reduces the number of unique equations from (*N* + 1)(*N* + 2)*/*2 to *N* + 1.

**Figure 2:**
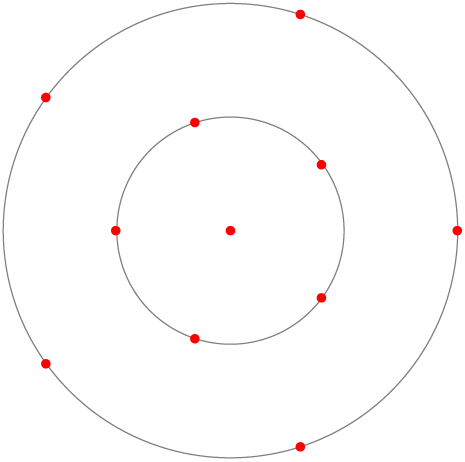
An example dipole placement with one dipole at the origin, and two rings of five dipoles. The outermost ring can have two possible orientations; the shown one, i.e., not aligned with the innermost ring, can solve all moments up to the degree of 6.

## 3. Methods and models

We developed and evaluated our methods using computer simulations in a realistic head model. First, we evaluated the coil-related approximations and built and tested real-time solver implementations using our previously-verified [20, 21] solver and four-compartment head model that contained brain + cerebellum, CSF, skull and scalp compartments. Then, we implemented five-compartment models that contained separate gray and white brain matter, evaluated discretization errors and performed model comparisons and speed evaluations.

All computations were done using Linux servers and workstations. For part of speed testing, we used C language (gcc 5.4.0), all other computations were done in MATLAB (versions R2017b or newer, www.mathworks.com).

### 3.1. Coil models

Our coil model are based on the description of the Magstim 70mm Double Coil in [15]. For reference coil, we divided the example coil into 1 mm^3^ cubic voxels, each represented by one dipole. Current was assumed to be spread evenly within the wires, and the weight of the dipole was set to the mean value of *f* in the voxel. This discretization approach, illustrated in Figure 3, resulted in a total of 85288 dipoles in 7 uniformly spaced layers. In test coil models, we had 1–3 layers with 2–42 dipoles in each, resulting in altogether 27 test coils containing 2–126 dipoles. In addition, we tested the Thielscher–Kammer (TK) coil model [15] that has 2712 dipoles.

**Figure 3:**
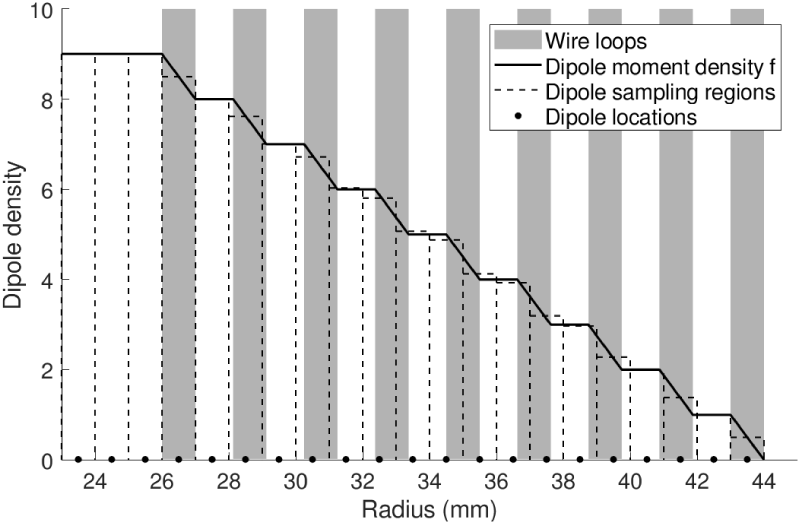
Discretization of our reference coil into a set of magnetic dipoles: The locations of wire loops are shown with gray rectangles, and dipole moment density f is plotted with a black line. The sampling regions and equivalent value of f for each dipole is shown with dashed line, and the dipole locations are shown with black dots.

### 3.2. Anatomical models

Our head models were built on the boundary meshes of the anatomical model of the sample data set of SimNIBS software (version 2.0)[22]. We resampled the meshes to various densities using iso2mesh toolbox [23] and in-house tools. Key information of our meshes is collected in Table 1.

**Table 1:** Anatomical models and the meshing densities of white–gray and pial surfaces. Models 5Cb25 and 5Cb30 have five conductivity compartments, models 4Cpxx have four. TSL marks the mean triangle-side length, N ^v^ the number of vertices. For the outer meshes, the 4Cref model had the TSLs of 3 mm for the cerebellum, 3 and 4 mm for the inner and outer skull, and 5 mm for the scalp. Other meshes had the corresponding TSLs of 5, 5, 6, and 7 mm.

| Name | gray-white | | pial | | Total $N^v$ |
| --- | --- | --- | --- | --- | --- |
| | TSL (mm) | $N^v$ | TSL (mm) | $N^v$ | |
| 5Cb25 | 2.5 | 27421 | 2.5 | 32496 | 69672 |
| 5Cb30 | 3.0 | 18424 | 3.0 | 21754 | 49933 |
| 4Cref |  |  | 2.0 | 52019 | 75652 |
| 4Cp20 |  |  | 2.0 | 52019 | 61774 |
| 4Cp25 |  |  | 2.5 | 32496 | 42251 |
| 4Cp30 |  |  | 3.0 | 21754 | 31509 |
| 4Cp40 |  |  | 4.0 | 11298 | 21053 |
| 3S |  |  |  |  | 8857 |
| 1S |  |  |  |  | 3510 |

For basic method development and coil model development and evaluation, we used a four-compartment (4-C) test model 4Cp40. This model is illustrated in Figure 4. For verification, we used a very dense 4-C reference model 4Cref and 4-C test models with varying TSLs in the pial mesh. Based on the results of this verification, we built five-compartment (5-C) models 5Cb25 and 5Cb30) that have the TSLs of 2.5 or 3 mm in both white–gray and pial boundaries and the same outer boundary meshes as the 4C test models. For comparison, we also built three-shell (3S; intracranial space, skull, scalp) and single-shell (1S; intracranial space) models. Models 4Cp40 and 4Cref are the same as Test 4 and Reference models of Head 1 in [21]; please see that publication for more detail on mesh processing.

**Figure 4:**
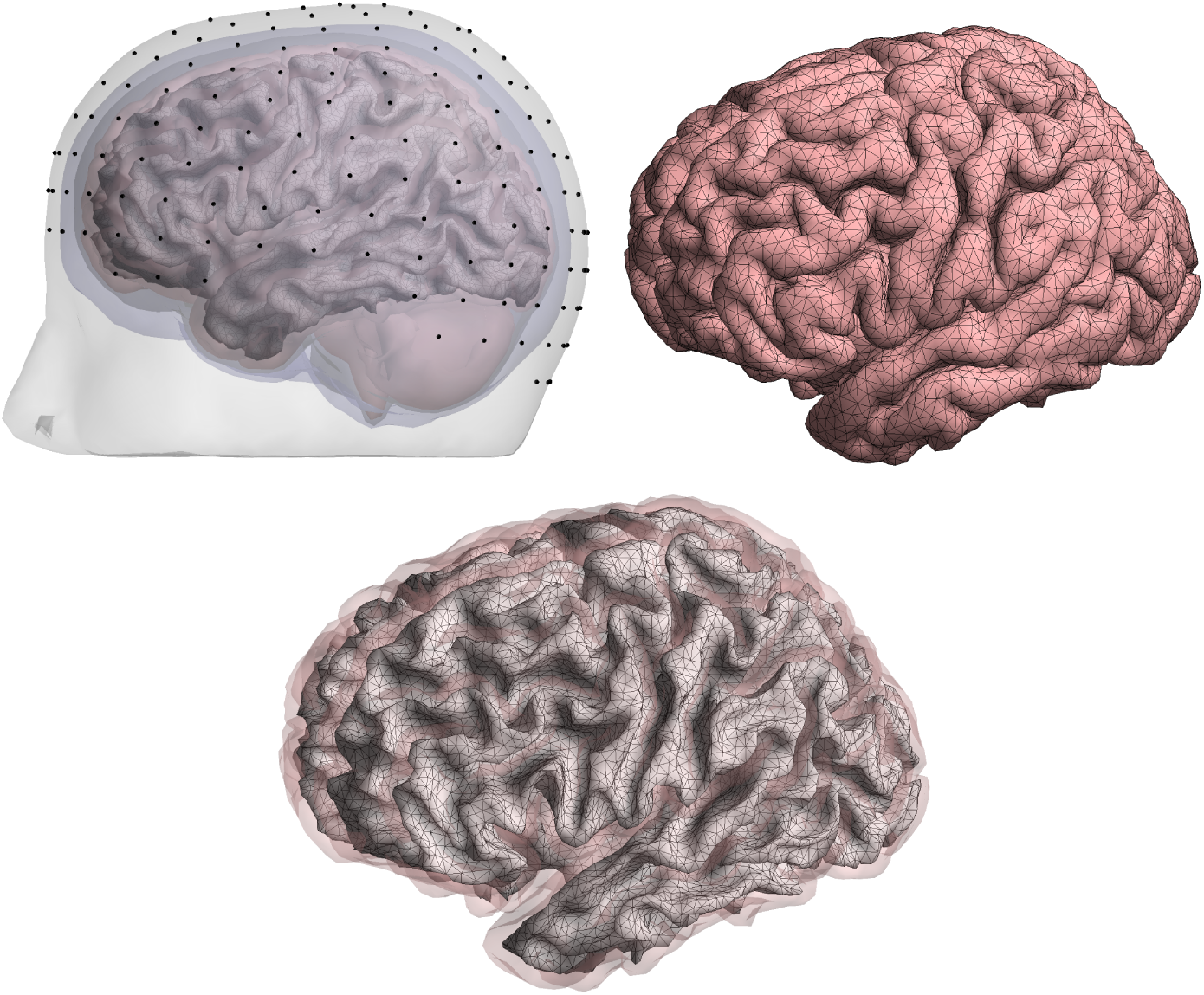
Computation geometry. The plot on top left shows all conductivity boundaries and also the field computation surface. The plot on top right shows the boundary between the brain and CSF, and the plot in the bottom shows the field computation surface in light gray and the pial surface in transparent light red. The locations of the stimulation coil (center of the coil projected to scalp) are shown with black dots in the top left-side plot, and the field computation points are in the vertices of the field computation surface mesh.

The E-field was computed in the cortex with 3-mm spacing, in vertices of meshes resampled from original FreeSurfer [24] meshes. In coil development and numerical verification using 4C models, the field was computed on both hemispheres. The field points (i.e. positions of dipoles 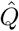) were placed on the boundary of the white and gray matter as in typical surface-based EEG and MEG applications, ensuring 1.5 mm distance between the dipoles and the pial mesh as in [21]. In all computations and comparisons that involve 5-C models, we computed the E-field in the left hemisphere. The field points were placed halfway between the white–gray and pial surfaces, first spanned on the resampled mid-surface of FreeSurfer brain surfaces and then projected so that each dipole was as far from both of our 2.5-mm brain meshes.

### 3.3. Boundary potentials

The model compartments were assigned conductivities of 0.14 S/m for white matter, 0.33 S/m for the gray matter, cerebellum and scalp, 1.79 S/m for the CSF, and 0.0066 S/m for the skull; these values are typical in EEG/MEG applications. The boundary potentials ***ϕ*** for all unit-current dipoles 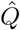 in xyz orientations were solved using boundary-element method formulated with isolated source approach, applying linear basis functions and Galerkin weighting (LGISA) [20]. The integrals of weighted dipole potentials in Eq. (11) of [20] were solved analytically as described in [25] instead of using numerical quadrature. This small change removes the well-known numerical stability issue related to placing dipoles near conductivity boundaries. For further detail, please see [20, 21] and Section 5.2.

### 3.4. Simulations

The coil was placed tangentially over the scalp in 251 separate positions around the head, following a 256-electrode EEG electrode layout. The coil was oriented so that the primary E-field would be approximately normal to the cortex in nearest sulci under the coil center. For each coil position, we computed the E-field across the whole cortex using different coil models and both accurate and approximated *β* integrals.

We compared the fields using relative error (RE), correlation error (CCE), relative magnitude (MAG), and mean angle error (AE) metrics. Let (*N* ^ROI^× vector arrays **E**^ref^, **E**^test^ contain the E-fields computed in reference and test models; their difference is **E**^diff^ = **E**^test^ − **E**^ref^. Arrays de-meaned across field points are marked with subscript c, vector arrays pooled into one (3*N* ^ROI^ 1) scalar array with subscript p, and the L2 norm with horizontal bars ‖. Our main metric is the relative error that measures the overall difference of the arrays,

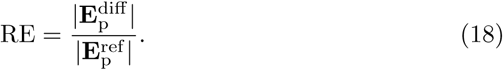

Correlation error characterizes the differences of the shapes of field patterns, ignoring constant amplitude differences:

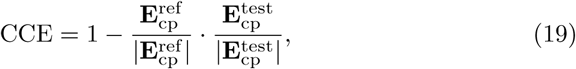

where the dot marks the scalar product and de-meaning is done before pooling. Relative magnitude simply compares the total amplitudes of fields,

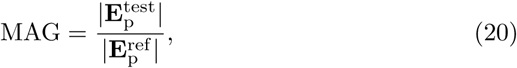

and mean angle error compares field orientations pointwise and takes mean across all field points. All metrics were computed both for all field points and thresholded to those points that had E-field energy density over 50% of the maximum (i.e. over 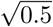 times the maximum E-field magnitude) for that coil location.

For selected coil models, we performed speed testing. In the first tests, we solved the normal component of 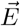 in either the whole brain (number of field points *N*^b^) or in a region-of-interest around the projected coil center. Potential ***ϕ*** was pre-computed for all *N*^b^ oriented sources on the cortex, resulting in (*N*^v^ × *N*^b^) matrix **Φ**. For each coil position, following computations were done:

1. Build **D** and compute **d** = **w**^T^**D**, resulting in a (1 × *N*^v^) vector.
2. Build **B**^p^ for all field points and compute **w**^T^**B**^p^, resulting in a (1 × *N* ^ROI^) vector.
3. Compute 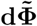, where 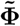 is either the full (*N*^v^ × *N*^b^) matrix **Φ** or *N*^ROI^columns of it.

In the second part, we solved the full 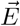 in a static ROI with varying number of points, using models with various sizes *N*^v^; 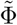 was then of size (*N*^v^ × 3*N*^ROI^). These tests were done using the GPU-accelerated MATLAB implementation.

The benchmarking computer had a 4-core processor (Intel Xeon E3-1230 v5, 3.4 GHz) and a consumer-level graphics card (Nvidia GeForce GTX 1060). The MATLAB implementation used the GPU via `gpuArray` functionality for computing the multiplication 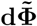, while the C implementation used only the CPU. The C implementation used single-instruction-multiple-data (SIMD) instructions supported by the processor when applicable, and both implementations used all CPU cores when applicable.

### 3.5. Data and code availability

To facilitate further development in the research community, our dipole-based coil models and example MATLAB routines for field computation are available from the corresponding author for academic use.

## 4. Results

### 4.1. Coil model parameters

We solved the z-coordinates of dipole layers for an origin-centered planar coil of thickness *d* as described in Eqs. 11–15. Eq. 15 was solved for one to four layers *I* using Mathematica software (version 10.3.0, www.wolfram.com). This resulted in optimal layer positions and weights shown in Table 2. The layers are placed less densely in the center and more densely towards the coil boundaries, while the central layers have higher weights than the ones closer to the boundaries.

**Table 2:** Optimal layers (z_i_) and their weights (w_i_) for discretisation of a planar coil with thickness d, centred around origin.

| $I$ | $z_i/d$ | | | | $w_i$ | | | |
| --- | --- | --- | --- | --- | --- | --- | --- | --- |
| 1 | 0 |  |  |  | 1 |  |  |  |
| 2 | -0.2887 | 0.2887 |  |  | 0.5 | 0.5 |  |  |
| 3 | -0.3873 | 0 | 0.3873 |  | 0.2778 | 0.4444 | 0.2778 |  |
| 4 | -0.4306 | -0.1700 | 0.1700 | 0.4306 | 0.1739 | 0.3261 | 0.3261 | 0.1739 |

Then, we sought the optimal distribution of dipoles, looking for the minimum number of dipoles that produces the exact match of moments 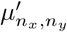 and 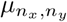 up to a degree *n*_*x*_ + *n*_*y*_ = *N* for a system of circular rings. A systematic search through all possible sets of dipoles for *N* ≤ 9 gave us the dipole placement rules shown in Table 3. Unlike with *z* direction, the weights and radii cannot be given in closed form for an arbitrary coil, so we continued by fitting the radii and weights for one wing of our example coil, where each wire loop (shown in Figure 3) was further represented with 10 equally spaced thin circular loops of Eq. 17, and the system of polynomial equations was solved using Mathematica. At the end we then had 9 sets of dipoles in the central plane of the coil. With these, we implemented the whole coil using 1, 2, or 3 layers, resulting in 27 test coil models.

**Table 3:** Dipole placement rules for minimum number of dipoles to match all moments up to a degree of N of a system of planar circular disks. The 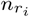 dipoles in each ring r_i_ have equal weights and equidistant spacing of 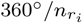, and the two rings of dipoles for the last four columns have an offset of 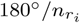. Subscript o indicates the origin and P P L shows the total number of dipoles per layer of the whole coil model. All shown cases have seemingly more equations (N + 1) than unknowns (n_o_ + 2 · n_rings_); after simplification, each case has, however, exactly n_o_ + 2 · n_rings_ unique equations.

| $N$ | 1 | 2 | 3 | 4 | 5 | 6 | 7 | 8 | 9 |
| --- | --- | --- | --- | --- | --- | --- | --- | --- | --- |
| $n_o$ | 1 | | | 1 | 1 | 1 | 1 | 1 | 1 |
| $n_{r_1}$ | | 3 | 4 | 5 | 6 | 5 | 6 | 9 | 10 |
| $n_{r_2}$ | | | | | | 5 | 6 | 9 | 10 |
| PPL | 2 | 6 | 8 | 12 | 14 | 22 | 26 | 38 | 42 |

### 4.2. Coil model comparisons

We evaluated the accuracy of the *β* integral and coil approximations in model 4Cp40 by comparing the cortical E-fields of each test coil to those obtained with the reference coil model (85288 dipoles) and accurate *β* integrals. For all test coil models and accurate and approximate *β*, we computed the RE, CCE, MAG and AE both in all field points and thresholded to the region of the strongest field. REs for all field points across all test coils are shown in Fig. 5. We see immediately that single-layer coils perform better than the multi-layer coils with respect to the total number of points. All coils with at least 38 points per layer (PPL) have the mean RE below 1%. With PPL ≥12 we reach RE < 3%, while with PPL < 12 the error increases quickly. The 38- and 42-dipole coils perform as well with accurate *β* integrals, but with approximate integrals the 42-dipole coil has clearly smaller standard deviation. Other metrics did not bring relevant additional information.

**Figure 5:**
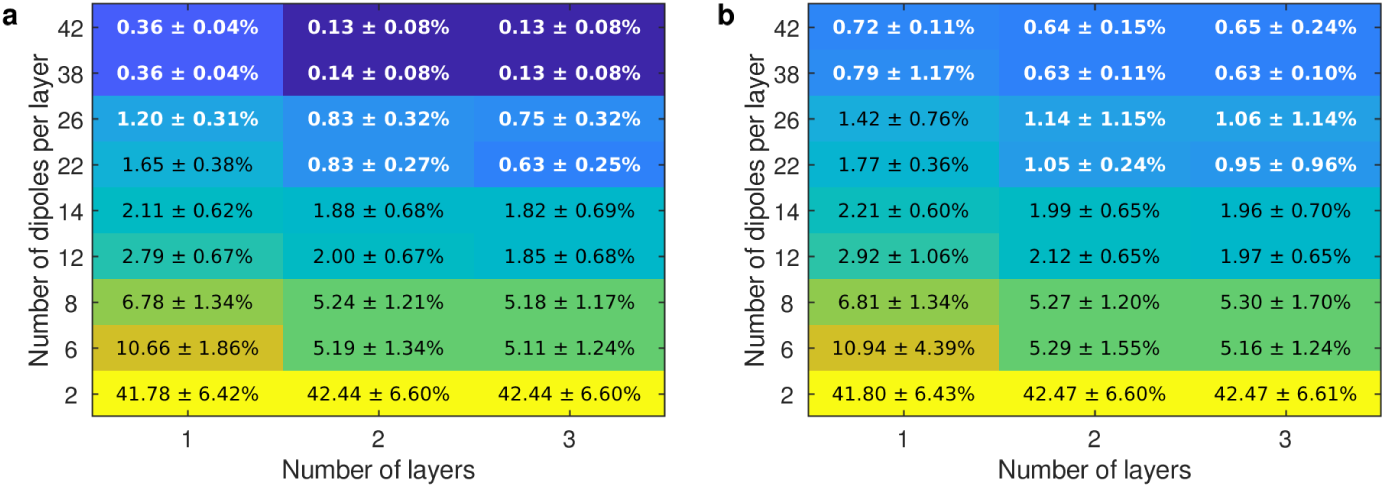
Relative errors due to coil model simplification and fast numerical integration of secondary magnetic fields, mean ± standard deviation across all 251 coil locations: a) test coils with accurate β integrals; b) test coils with fast numerical β integrals. The values printed in white are less than twice the error of the fast numerical approximation of β integrals in the reference coil.

In regions of strongest fields, the REs are for coils with PPL ≥ 38 slightly smaller than the total REs, and for coils with PPL between 12 and 26 slightly larger. The mean RE due to the fast numerical approximation of *β* integrals (Eq. 8) in the reference coil was 0.60 ± 0.09%. With test coils, the difference between the accurate and approximate integrals was smaller, as seen from Fig. To compare to previously published coil model [15], the TK coil model with 2712 dipoles had the RE of 4.1 ± 0.1% with both accurate and approximate *β* integrals.

To see, how much the mesh density affects the coil accuracy, we solved the E-field for 10 coil positions using the 4Cref meshing (75652 vertices) and both the reference coil (using accurate *β* integrals) and the 42-dipole coil. The mean ± standard deviation of the RE due to the 42-dipole coil was in all field points 0.41 ± 0.06% and in the region of strong E-field 0.38±0.03%; corresponding numbers in the coarser meshing 4Cp40 (21053 vertices) were identically 0.41±0.06% and 0.38±0.03%. When the test coil model used approximate *β* integrals, the errors with 4Cref mesh were 0.44±0.06% and 0.45±0.05% and with the 4Cp40 meshing 0.68±0.05% and 0.63±0.10%. The error of the coil model is thus not sensitive to the mesh density, and in denser meshes the *β* approximation increases the error less than in coarser meshes.

### 4.3. Five-compartment model and model comparisons

Based on the results of the multi-resolution verification presented in Appendix A, we built 5-C models 5Cb25 and 5Cb30. For all 5-C, 4-C, 3S and 1S models, we computed the full E-field in the mid-surface of the left-hemisphere cortex using the 42-dipole coil and those 105 coil locations that had the regions of the strongest E-field in the left hemisphere. Then, we compared the fields, using the 5Cb25 model as reference. Results with approximate *β* integrals are presented in Table 4; corresponding results for accurate *β* integrals are shown in Table C.7. Example E-fields on the mid-cortical surface are visualized in Fig. 6, and an example of high-resolution computation in a volume slice is shown in Fig. B.8.

**Table 4:** Discretization, model simplification and β approximation errors in 5-C and 4-C models meshed at various densities and solved with approximate β integrals. The dense 5-C model (5Cb25) with accurate β integrals is used as the reference, and the field space is in the mid-surface between the white–gray and pial boundaries. For each model, the upper row shows the error metric computed for all field points, and the lower row shows it for the region of strongest E-field. The results are presented as mean ± standard deviation. For corresponding results with approximate β integrals, see Table C.7.

|  | RE (%) | CCE (%) | AE (deg) | rMAG (%) |
| --- | --- | --- | --- | --- |
| 5Cb25 | $0.6 \pm 0.1$ | $0.0 \pm 0.0$ | $0.2 \pm 0.0$ | $100.0 \pm 0.1$ |
| | $0.4 \pm 0.2$ | $0.0 \pm 0.0$ | $0.1 \pm 0.1$ | $99.8 \pm 0.1$ |
| 5Cb30 | $5.2 \pm 0.6$ | $0.1 \pm 0.0$ | $1.8 \pm 0.0$ | $100.3 \pm 0.1$ |
| | $4.2 \pm 1.5$ | $1.9 \pm 1.5$ | $1.3 \pm 0.2$ | $100.1 \pm 0.5$ |
| 4Cp20 | $19.6 \pm 0.8$ | $1.5 \pm 0.1$ | $6.9 \pm 0.3$ | $109.1 \pm 1.0$ |
| | $11.7 \pm 1.6$ | $11.2 \pm 4.4$ | $3.6 \pm 0.5$ | $101.5 \pm 1.3$ |
| 4Cp25 | $19.0 \pm 0.8$ | $1.4 \pm 0.1$ | $6.8 \pm 0.4$ | $108.8 \pm 1.1$ |
| | $10.8 \pm 1.7$ | $10.0 \pm 4.3$ | $3.6 \pm 0.5$ | $101.2 \pm 1.3$ |
| 4Cp30 | $19.2 \pm 0.8$ | $1.5 \pm 0.1$ | $7.0 \pm 0.3$ | $108.6 \pm 1.1$ |
| | $11.2 \pm 1.6$ | $10.9 \pm 4.6$ | $4.0 \pm 0.5$ | $101.0 \pm 1.4$ |
| 4Cp40 | $20.1 \pm 0.7$ | $1.7 \pm 0.1$ | $7.7 \pm 0.4$ | $108.0 \pm 1.3$ |
| | $12.4 \pm 1.6$ | $13.5 \pm 5.7$ | $4.8 \pm 0.6$ | $100.6 \pm 1.7$ |
| 3S | $27.1 \pm 1.0$ | $4.0 \pm 0.3$ | $12.1 \pm 0.7$ | $97.2 \pm 1.0$ |
| | $22.5 \pm 2.4$ | $23.5 \pm 6.9$ | $6.7 \pm 0.9$ | $87.0 \pm 3.0$ |
| 1S | $28.0 \pm 1.4$ | $4.2 \pm 0.4$ | $13.5 \pm 1.2$ | $95.0 \pm 1.9$ |
| | $23.7 \pm 2.7$ | $23.6 \pm 7.4$ | $6.7 \pm 0.9$ | $84.8 \pm 3.6$ |

**Figure 6:**
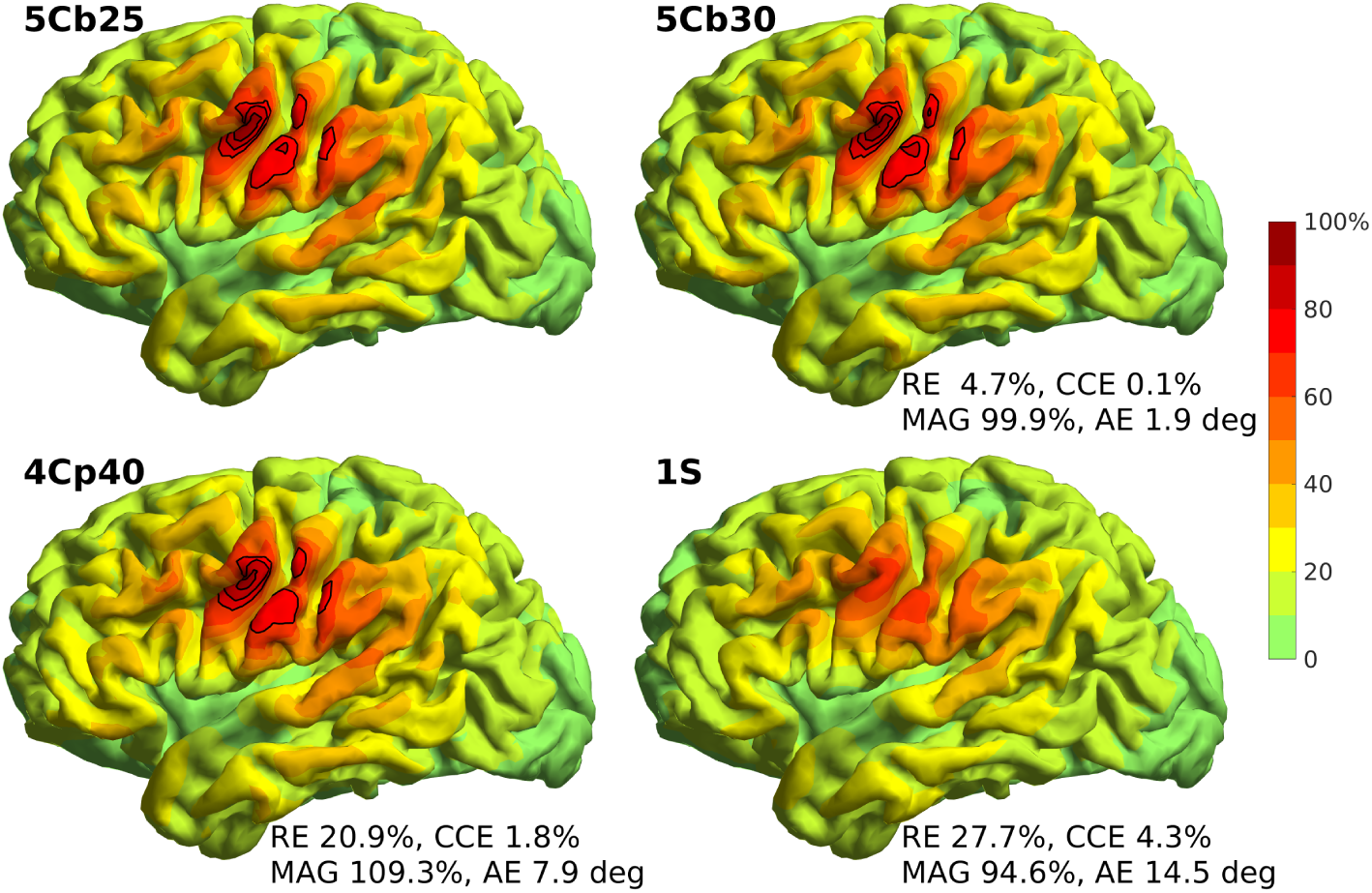
Example E-field distributions and error metrics for one coil position. The 5Cb25 model (top left) with the reference coil and accurate β integrals was used as reference, while the test models 5Cb30, 4Cp40 and 1S used the 42-dipole coil and approximate β integrals. The E-field was computed on the mid-surface of the cortex. The colormap codes the magnitude of the E-field in percents with respect to the maximum of the reference data, and the contour lines depict the field over 70% of that maximum.

The RE and MAG of the 4-C models are 19–20% and 108–109% for all field points and 11–12% and 101–102% for the regions with the strongest field, while the correponding CCEs are 1.4–1.7% and 10.0–13.5%. This means that the 4-C model produces relatively larger field outside the region of the strongest field, while in the strongest region the overall field magnitudes match well. The field shape, on the other hand, has smaller differences in weaker regions than in stronger regions. The difference between errors of dense and coarse 4-C models is of the order of 1%-unit. The coarser 5-C model has 4–5% RE due to the discretization of anatomy and potential, while the 3S and 1S models have overall REs between 27–28% for all field points and 23–24% in the region of strongest field.

To compare the discretization error (4–5%) to other expected errors, we varied the conductivity of the brain compartments of the 5Cb30 model by 20% and solved the E-fields. When the conductivity of white matter was 20% smaller than expected, the mean RE for all field points was 4.7%; 20% larger conductivity produced a mean RE of 3.9%. Corresponding errors in gray matter conductivity produced REs of 2.9% and 2.4%, and if both conductivities were 20% off, the mean REs were between 4.4% and 5.4%. In the regions of strongest E-field, the mean REs were 1.6–3.5%. Similar results were obtained in [26]. The discretization error and conductivity error are thus of the same order.

### 4.4. Speed tests

We chose the 42- and 14-point single-layer coils and the 76-point two-layer coil for optimization and testing the speed of our different implementations. In MATLAB and C implementations and with the 42-dipole coil, the approximate *β* integrals were approximately 63 and 45 times faster than the accurate integrals. Thus we continued the speed tests with approximate *β* integrals only.

First, we computed in the 4Cp40 model (*N*^v^ = 21052) the normal component of E-field for either the whole cortex (*N*^b^ = 20324) or a region-of-interest (ROI) (mean *N*^ROI^ = 1805, cortical surface area 117 cm^2^, radius approximately 6 cm, each coil location having its own ROI). The computations were done sequentially for all 251 coil locations and repeated many times.

The total computation speeds field for approximate *β* integrals and different implementations are presented in Table 5. With the 42-dipole coil, we were able to compute the normal E field on all 20324 points of field space at the speed of 82.9 coil positions per second (cps). Computing all three field components took about three times as long (for example, full E-field in the whole brain 27.3 and 22.8 cps for 42- and 76-dipole coils in MATLAB with GPU acceleration).

**Table 5:** Computation speed with different coil models and code implementations, presented as coil positions per second. The computations for the whole brain had N^b^ = 20324 field points and those for regions of interest on average N^b^ = 1805 points.The size of the model was N^v^ = 21052 points.

| $N^c$ | Whole brain | | | ROI | | |
| --- | --- | --- | --- | --- | --- | --- |
|  | 76 | 42 | 14 | 76 | 42 | 14 |
| MATLAB (CPU) | 9.4 | 11.0 | 11.5 | 14.0 | 14.0 | 15.5 |
| C (CPU) | 13.2 | 13.6 | 13.9 | 107.5 | 114.1 | 123.1 |
| MATLAB (CPU+GPU) | 58.8 | 82.9 | 84.4 | 69.7 | 110.1 | 144 |

Next, using MATLAB+GPU and the 5Cb30 model with the 42-dipole coil, we studied, how the computation speed depends on model size. We computed all components of E-field varying either the number of field points *N* ^ROI^ or the size of the volume conductor model (i.e. *N* ^v^), resulting in 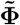 matrix with dimensions (*N* ^v^× 3*N* ^ROI^) and thus altogether 3*N* ^ROI^*N* ^v^ entries. The computations were done for all coil locations and repeated many times. The full model had 9000 field points and 49933 model vertices, the size being constrained by the 6 GB memory of our GPU. The results are visualized in Figure 7. For *N* ^ROI^ = 9000, the model was solved at the speed of 26.6 cps and for *N* ^ROI^ = 4000 at 64.3 cps. The computation time depends linearly on the system size, and the increase of computation time is mostly due to the increasing size of matrix **Φ**—the time needed for other computations grows much more slowly.

**Figure 7:**
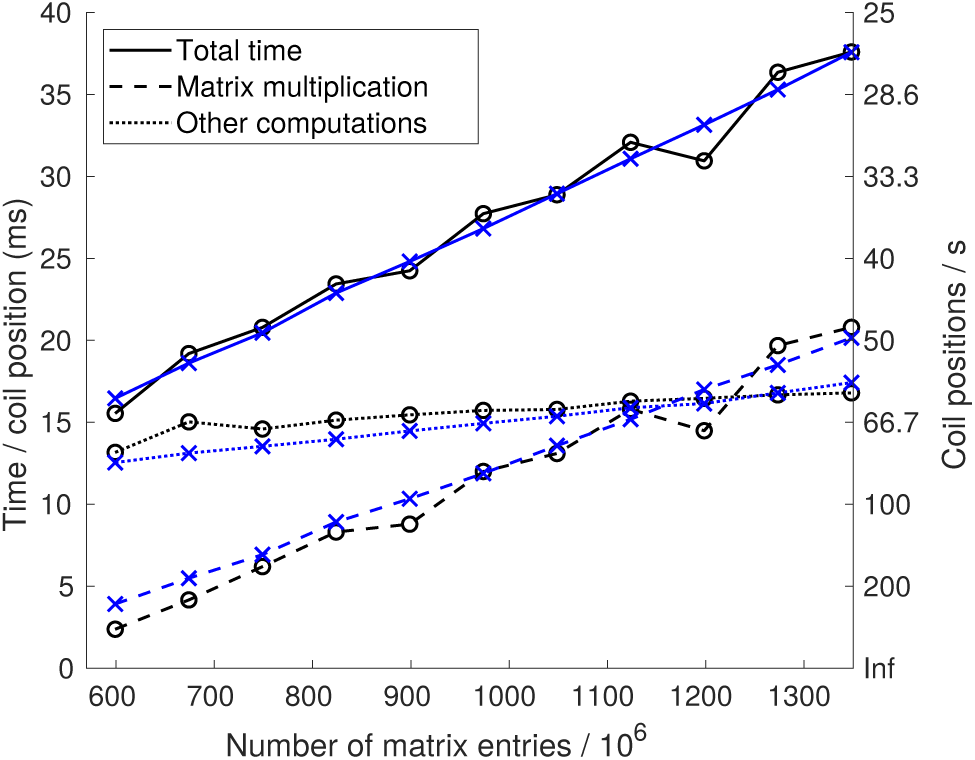
Computation time and speed per coil position as function of model size. The black plots with circular markers show the computation time for a model with 49933 vertices and varying number of field points, and the blue plots with cross markers correspond to a model with 9000 field points and varying number of vertices; the largest model has 49933 ·9000 ·3 = 1348 ·10^6^ matrix entries. The total time is plotted with solid line, the time spent in vector– matrix multiplication 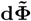 with dashed line, and the time spent in rest of the computations (β integrals, primary magnetic fields, coil mapping, loading the data to the GPU) in dotted line. The left y-axis shows the time per coil position, and the right y-axis shows the corresponding speed as coil positions per second.

Solving only the normal component in 9000 points of the 5Cb30 model ran at the speed of 81.0 cps i.e. three times the speed of the full field. We also tested models 4Cp30 and 4Cp40 with 9000 field points; all field components were solved at speeds of 42.7 and 64.0 cps and the normal component at speeds of 127.9 and 167.9 cps, respectively. With the dense 5Cb25 model and all components of the E-field, we reached real-time performance with 6500 field points (26 cps).

For reference, the offline part i.e. building **Φ** for 20000 source triplets took for the 4Cp40 model less than one hour on the benchmarking PC (CPU only), and for the 5Cb30 model approximately two hours on 12 cores of a standard server computer (Xeon E5 2680 v3). The majority of time goes into building BEM geometry matrices—a task that is done only once per meshing and that can be parallelized further, if faster offline computation is needed.

## 5. Discussion

In this paper, we presented methods for computing the TMS-induced E-field in the brain in real time. The real-time performance is reached with the help of 1) formulation that allows de-coupling of coil-related calculations from other calculations; 2) an approximated solution of the *β* integral used for computing the contribution of head conductivity on E-field; and 3) a new moment-based approach for approximating the TMS coil model.

We verified and demonstrated the methods and compared models using over 100 TMS coil locations and realistic four- and five-compartment (4-C, 5-C) volume conductor models (VCM) that contain (white and gray) brain tissue, cerebrospinal fluid, skull, and scalp. Using our dense 5-C model as reference and looking at the regions of strongest E-fields, the RE of our real-time 5-C model (5Cb30) was 4.2%, while the corresponding errors of medium-density and coarse 4-C models were 11.2% and 12.4%. The three- and single-shell models had the mean REs of 22.5% and 23.7%.

The error introduced by the approximated *β* integrals was between 0.3% and 0.6% depending on mesh density; coarses meshes produced larger errors. The relative error (RE) due to approximating the coil with 42 dipoles was approximately 0.4%. Together, the REs due to the new approximations were between 0.44% and 0.72%. When also other modeling errors were present, the contribution of these approximations to the total error was of the order of 0.1%- units. The error due to the approximations that enable real-time computation is negligible compared to the difference between models with different levels of detail and much smaller than typical mesh-related discretization error or errors due to unknown conductivity parameters.

On a standard PC with a consumer-grade GPU, the computation speed for all components of the E-field in 9000 field points was 27 cps in a 5-C model and 43 and 65 cps in medium-density and coarse 4-C models. Solving only the normal component of the E-field approximately tripled the speed. These speeds are clearly adequate for experimental TMS navigation use.

### 5.1. Coil model

We used Magstim 70 mm figure-of-eight coil as our example coil and modeled it following the specifications reported in [15]. A real-world coil has spiral windings, but we modelled the coil windings as a system of circular loops. This common simplification, used e.g. in [15, 2, 27, 6, 9] enables the use of analytically-computed reference moments, facilitating our method development considerably by removing one degree of numerical uncertainty. We solved also the dipole radii and weights analytically. Our dipole placement rules in Table 3 were derived for a system of circular disks, so they hold for all planar coils with circular loops; the radii of dipole wings and weights of dipoles will need to be fitted separately for different windings.

The optimization of our dipole placements is based purely on geometrical moments. The independency of the coil model parameters of E-field models should increase the robustness of the coil model. This approach does, however, not guarantee optimal approximation of the E-field. In addition, we considered only those moments that we expected to fit perfectly and ignored higher moments. But, as our simulations show, the moment-optimized coils perform very well in realistic field computations. Our coil models should also work with direct surface- and volume-based approaches. With volume-based approaches, one may, however, need a higher-order model with higher number of dipoles, because the field produced by a sparse set of dipoles may not be sufficiently smooth in the elements nearest to the dipoles.

The moment approach as presented here should work with all circular coils and figure-of-eight coils with circular wings. Most commercial coils used for navigated TMS and overall for TMS mapping are of this type. The approach can be generalized to non-planar and non-circular coils, but then one likely needs to resort to numerical computation of moments and thus has to pay attention to the numerical stability by, e.g., regularizing the fitting, especially in solving the weight equations. An alternative to numerical moment computation would be the combination of moments based on circular rings, geometrical morphing, and further optimization using E-field models. Our figure-of-eight coil has a high degree of symmetry, enabling excellent accuracy with a small number of dipoles. With a non-symmetric coil, for example one that has different shape for each wire loop, one may need to either accept a larger coil-related error or use a considerably larger number of dipoles and thus have smaller ROI or slower computation. We leave modeling of non-circular coils for further research.

Our 85288-dipole reference coil model was verified by comparing the magnetic vector potential 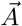 produced by it to an 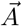 computed analytically in a line-segment model of the same coil (constructed independently). This comparison is equivalent to comparing primary electric fields. In the line-segment model, each circular loop was presented using 200 straight line segments (not split to dipoles), and the wire height and thickness were taken into account by dividing the cross-section of the wire evenly into 28 elements, each represented by one wire. For a coil above the primary motor cortex and field computed on the cortex, the difference (RE) between our reference coil and the line-segment coil was *<*0.008%. We also verified that our coils and algorithms produce the same primary E-field using both reciprocal and direct approaches.

Compared to our coils, the TK coil model with 2712 dipoles had a relatively large RE of 4.1*±* 0.1%. The reason for this is the weighting rule that leads to too large weights for dipoles positioned within wire loops: in the model description [15], those dipoles were given the full weight of the loop in question, although they should get only part of that weight (see the dipole moment density *f* and the dipole values of our reference coil in Figure 3). Using the dipole locations of the TK model and our weighting rule, we built a corrected TK model (TKc) that had the RE of 0.08 *±* 0.01%—a value expected of a coil with such a large number of dipoles.

### 5.2. Computational approach

Our computational approach is based on the principle of reciprocity and computing the secondary electric field by weighted surface integration of discretized boundary potentials ***ϕ***, while typical high-detail E-field models apply direct approach and volume discretization [2, 4, 28, 8]. The presented methodology does not depend on the way how the ***ϕ*** is solved—even though we used a boundary-element method [20], one can also use finite-element or finite-difference methods. We used the same mesh in computing the potential and the *β* integrals; if one uses different meshings or basis functions in these steps, one has to pay attention to not corrupt the calculation by interpolation errors.

The main benefit of the our approach, partially enabling the high computation speed, is that the computations specific to the coil can be efficiently separated from those that characterize the overall volume-conductor behavior. Related to that, the point set or mesh, where the E-field is computed, is independent of the discretization of the volume conductor model. Thus, the density of E-field sampling and speed of computation can be easily varied—for example, we could apply a coarse mesh in the first stages of navigation, and change to a more detailed one in later stages of navigation that demand higher sampling density or more accurate visualization. A further benefit is that the computation is done directly in terms of electric field instead of computing first the potential and then obtaining the (secondary) field via numerical differentiation over elements, as is typically done with direct volume-based approaches [2, 4, 28].

The key limitation is that the Geselowitz surface integral equation, Eq. 3, is valid only for isotropic medium; thus, the anisotropic conductivity of white matter cannot properly be accounted for. According to FEM simulations [27], the anisotrophy has a very small effect on E-field amplitude in gray matter, but in white matter the effect is on average 7% (max. 40%). Thus, if one aims to model the E-field in white matter, imaging data needed for modeling the anisotrophy is available, and real-time performance is not required, one might want to choose the direct approach and FEM modeling instead of the reciprocal approach as used here.

The main computational challenge in implementing the real-time reciprocal approach is that we need to place dipoles in all field points and pre-compute their potentials; in a dense (volume) grid, the number of these dipoles and thus the size of the **Φ** matrix would increase quickly, making the problem too large to tackle, especially on a GPU. The real-time implementation is thus best suited for computing field on a surface in the brain instead of volume grids. The size of the problem is also directly proportional to the number of vertices *N* ^v^ in the conductivity boundaries. Thus, if adding more model compartments, it makes sense to optimize the meshing—a geometrically simple boundary not directly adjacent to the cortex can typically be represented with a rather coarse mesh.

One also has to pay attention on the numerical stability of the solution of the electric potential, when field points are very close to a conductivity boundary (say, *<* TSL/2); such situations occur regularly, when meshing the cortex boundaries with elements whose TSLs are of the order of cortex thickness. In Galerkin-weighted boundary element methods, this problem is mainly due to the interaction between the pointlike sources and quadrature points: when a source is very close to a boundary element, a small shift in the source position may cause a large difference in distance to the nearest quadrature point, leading to a large change in the weighted dipole potentials (Eq. (11) of [20]). In our implementation, this numerical issue was cured by solving the said integrals analytically.

In addition to our reciprocal approach, the surface-integral solution for TMS-induced E-field can be formulated in direct way via induced surface charges [9, 10, 29]. Both approaches follow from quasi-static Maxwell equations without further approximations and are hence physically equivalent. In associated integral equations, the distance-dependent terms are of the same order, leading to similar effects of discretization errors. With the same meshing and matching discretization and integration techniques, the E-fields should thus be nearly identical, apart from small numerical differences. We implemented the direct approach following [9, 29], using the same linear discretization and integration techniques as in our approach, and computed the cortical E-field for 251 coil positions in a 4-C model. The mean relative difference between the approaches was 0.28%. Omitting the isolated-source formulation ISA (that in general improves the numerical behavior) in the reciprocal approach, the difference dropped to 0.006%.

The BEM-FMM in [9, 10, 29] is a direct BEM optimized for solving dense models. In this optimization, field integrals that we solve accurately are approximated using single-point quadrature and fast multipole method (FMM), and the linear equation system is solved approximately. Due to these choices and not using ISA, the BEM-FMM is expected to be numerically slightly less accurate than our approach with fast *β* quadrature. This lower intrinsic accuracy is compensated by higher mesh density. Comparing BEM and FEM with corresponding boundary geometries and element orders, BEM is expected to have higher accuracy [10, 29, 30]; to reach the same numerical accuracy, FEM thus needs a considerably denser mesh or higher element order. With the FEM—especially the hexahedral FEM of [5]—it is, however, relatively easier to increase the mesh density and thus improve the accuracy of both numerics and geometry. Interpreting results of our verifications and recent simulations [29, 31], our discretization error of 4–5% is likely dominated by the geometry error. Concerning the intended use of our real-time approach and being aware of differences, approximations and errors in MR images [32], segmentation [33] and conductivities [26], the overall discretization error of our real-time 3-mm model should be reasonably close to the corresponding errors of the high-density BEM-FMM model of [9] and the default SimNIBS 3.0 model of [31].

The added error due to ignoring the separation of white and gray matter is of the same order with some model differences presented in [10, 29, 26] and smaller than the approximation errors in the fast solution of [8]. That said, the choice between a 5-C and 4-C model is not just a question of accuracy under ideal simulation conditions: taking an experimental point of view and admitting the errors mentioned in the previous paragraph, the 4-C model provides an accessible solution for experimenters, who wish to upgrade from spherical models but have no resources or skills for making fast 5-C models.

### 5.3. Computation speed

With the formulation, approximate *β* integral and coil models presented in this work, we can in a standard desktop PC equipped with consumer-grade GPU solve the full E-field in 9000 field points on the cortex in real time using a detailed realistic head model; depending on the level of detail and meshing of the head model, the computation speed was between 27 and 65 coil positions per second. Solving only the normal component of the field approximately triples the speed. The computation speeds depends essentially only on the size of the model, i.e., the number of field points *N* ^ROI^ times the number of vertices *N* ^v^ in the conductivity boundaries of the head model—with a 1800-point ROI and 4-C model, the real-time performance can be reached even without the GPU. In the examples of this work, the model size is constrained by the 6 GB memory of our (low-spec) GPU. With higher-spec GPU, we can further increase either the size of the model, the computation speed, or both.

The computational load is dominated by the matrix–vector multiplication **dΦ**. In CPU-only computations with a coarse 4-C model and the whole brain, the C program was somewhat faster than the MATLAB one. The GPU implementation of **dΦ** speeded the MATLAB computations up by a factor of six. Assuming the same GPU libraries, a C+GPU implementation would with large models likely run only slightly faster than the MATLAB+GPU implementation, the benefit coming from faster building of arrays **d** and **B**^p^. Here, we used the GPU only for computing **dΦ**, using the simple gpuArray functionality in MATLAB. With more invoved GPU programming, one could build **d** and **B**^p^ on the GPU and potentially further cut down the computation time.

When computing in a coil-specific ROI, the C version performed about six times as fast as in the whole brain, while the MATLAB CPU and CPU+GPU programs got speed gains of only 40–50% and 18–30%, respectively. The main reason for this difference lies in the implementation of matrix–vector computation: Both CPU and GPU libraries used by MATLAB slow down considerably when they operate on only part of the columns of **Φ**. Also the memory consumption increased, hinting that the libraries make copies of the needed columns instead of operating directly on them. Thus, in further tests with the GPU, we used a large static ROI.

Our static ROI was had the size of 9000 field points. With our 5-C model, the number of entries in **Φ** matrix was then 3 *·N* ^v^ *·N* ^ROI^ = 3 *·*49933 *·*9000 = 1.348 *·*10^9^, requiring 5.02 GB of memory with single-precision floats. This size is enough to cover, for example, cortical regions of one hemisphere meshed at 3 mm density. Loading a new such ROI onto the GPU takes less than one second. We leave further optimization and study of GPU acceleration for future research.

### 5.4. Outlook and future research

Here we have presented, verified and benchmarked new computational methods that enable real-time E-field computation in realistic head geometry that includes realistic CSF and gyral structure and isotropic white matter. The new methods should have impact on all TMS applications, where more accurate targeting and precise dosing of stimulation during the TMS experiment or treatment is desired.

In near future, we plan to extend our coil modeling to other coil types and to overall develop and optimize our solver implementation. Further work is also needed in generation and optimization of anatomical models and meshing.

## Acknowledgements

Computational and storage resources of Aalto University School of Science Science-IT project are acknowledged.

## Appendix A. Multi-resolution verification of the 4-C model

To assess the effect of discretization errors on the E-field in the 4-C model and the error due to *β* approximation, and to select meshing parameters for the 5-C models, we compared the dense 4Cref model to our other 4C models. The results computed for the 42-dipole coil in 251 locations are shown in Table A.6. In regions of strong E-field, the computationally light 4Cp40 model has an overall mean RE of 5.4% and mean CCE of 3.2%; the errors decrease clearly, when changing to the 4Cp30 model that has the TSL of 3 mm in the pial surface. Comparing Table A.6a with Table A.6b, we see that with approximate *β* integrals the RR, CCE and AE metrics and MAG means increase at most 0.1 %-units, while the MAG standard deviations vary up to 0.8 %-unit.

We also applied the reference coil model (in 10 locations) in the reference computations. Compared to results obtained with the 42-dipole coil as reference, the RE, CCE and AE values varied up to 0.1%-units or degrees, the MAG values up to 0.4%-units. Taken together, the increase of error due to *β* approximation and 42-dipole coil is small and in practice negligible, when any other model errors are taken into account. In further model comparisons involving the 5-C model, we can thus safely use the 42-dipole coil model as reference.

Model 4Cp20 has the same pial mesh as the reference model 4Cref and the same outer meshes as models 4Cp25, 4Cp30 and 4Cp40. The small errors of 4Cp20 show that the E-field problem is rather insensitive to the density of outer meshes; in 5-C models we can thus use the same coarse outer meshes as in the 4-C test models.

**Table A.6:** Discretization errors in 4-C models meshed at various densities and solved with accurate and approximate β integrals. The very dense 4-C model (4Cref) with accurate β integrals is used as reference, and the field space is at the gray–white boundary. For each model, the upper row shows the error metric computed for all field points, and the lower row shows it for the region of strongest E-field.

|  | RE (%) | CCE (%) | AE (deg) | rMAG (%) |
| --- | --- | --- | --- | --- |
| 4Cp20 | $0.8 \pm 0.3$ | $0.0 \pm 0.0$ | $0.3 \pm 0.0$ | $100.2 \pm 0.2$ |
| | $0.7 \pm 0.6$ | $0.1 \pm 0.1$ | $0.2 \pm 0.2$ | $100.3 \pm 0.5$ |
| 4Cp25 | $2.0 \pm 0.2$ | $0.0 \pm 0.0$ | $0.7 \pm 0.0$ | $99.9 \pm 0.2$ |
| | $1.8 \pm 0.6$ | $0.4 \pm 0.3$ | $0.6 \pm 0.2$ | $100.1 \pm 0.5$ |
| 4Cp30 | $3.4 \pm 0.2$ | $0.1 \pm 0.0$ | $1.2 \pm 0.0$ | $99.7 \pm 0.2$ |
| | $3.0 \pm 0.7$ | $1.0 \pm 0.6$ | $1.1 \pm 0.2$ | $99.9 \pm 0.7$ |
| 4Cp40 | $6.0 \pm 0.4$ | $0.2 \pm 0.0$ | $2.2 \pm 0.0$ | $99.1 \pm 0.2$ |
| | $5.4 \pm 1.2$ | $3.2 \pm 1.8$ | $2.0 \pm 0.4$ | $99.4 \pm 1.0$ |

b) Approximate $\beta$ integrals
|  | RE (%) | CCE (%) | AE (deg) | rMAG (%) |
| --- | --- | --- | --- | --- |
| 4Cref | $0.2 \pm 0.4$ | $0.0 \pm 0.0$ | $0.1 \pm 0.4$ | $100.0 \pm 0.0$ |
| | $0.2 \pm 0.1$ | $0.0 \pm 0.0$ | $0.1 \pm 0.1$ | $99.9 \pm 0.1$ |
| 4Cp20 | $0.9 \pm 0.3$ | $0.0 \pm 0.0$ | $0.3 \pm 0.1$ | $100.2 \pm 0.2$ |
| | $0.8 \pm 0.6$ | $0.1 \pm 0.1$ | $0.3 \pm 0.2$ | $100.1 \pm 0.4$ |
| 4Cp25 | $2.0 \pm 0.2$ | $0.0 \pm 0.0$ | $0.7 \pm 0.1$ | $99.9 \pm 0.2$ |
| | $1.8 \pm 0.6$ | $0.4 \pm 0.3$ | $0.6 \pm 0.2$ | $99.9 \pm 0.5$ |
| 4Cp30 | $3.4 \pm 0.2$ | $0.1 \pm 0.0$ | $1.2 \pm 0.1$ | $99.6 \pm 0.2$ |
| | $3.1 \pm 0.7$ | $1.1 \pm 0.6$ | $1.1 \pm 0.3$ | $99.6 \pm 0.7$ |
| 4Cp40 | $6.1 \pm 0.4$ | $0.2 \pm 0.0$ | $2.2 \pm 0.1$ | $99.1 \pm 0.2$ |
| | $5.5 \pm 1.2$ | $3.3 \pm 1.8$ | $2.0 \pm 0.4$ | $99.2 \pm 1.0$ |

## Appendix B. Example on high-resolution E-field computation

**Figure B.8:**
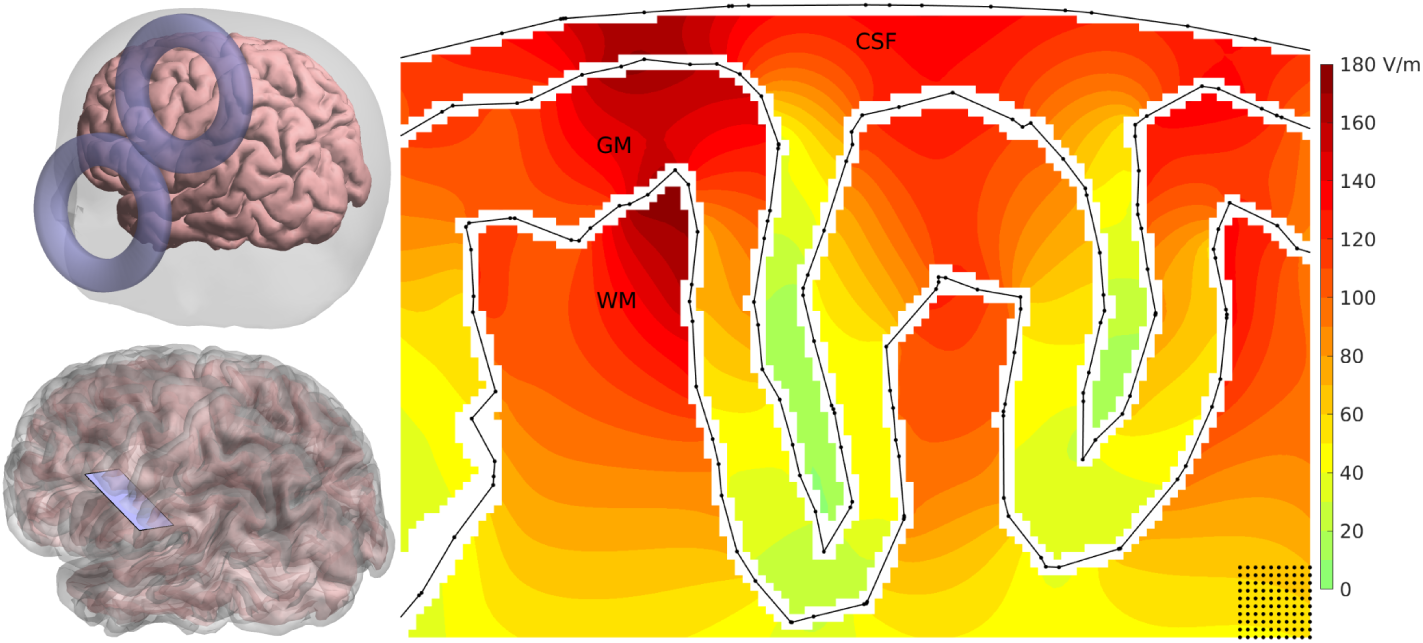
Example E-field distribution in volume, computed using the 5Cb30 volume conductor, 42-dipole coil model, and fast β quadrature. The field is computed under the coil center in a plane along the short axis of the coil as shown in the left-side plots. The isosurface plot shows the magnitude of the E-field (V/m) computed in a rectangular 0.3-mm grid omitting points that are less than 0.15 mm from the nearest boundary surface. Sample grid points are shown in the bottom right corner of the plot. Black solid lines show the projection of compartment boundaries on the plane, with black dots marking element edges. The size of the plane is 35 × 21 mm, and the coil is above the centerline of the plane.

## Appendix C. Model comparison results with accurate *β* integrals

**Table C.7:** Discretization and model simplification errors in 5-C and 4-C models meshed at various densities and solved with accurate β integrals. The dense 5-C model (5Cb25) with accurate β integrals is used as the reference, and the field space is in the mid-surface between the white–gray and pial boundaries. For each model, the upper row shows the error metric computed for all field points, and the lower row shows it for the region of strongest E-field. The results are presented as mean ± standard deviation.

|  | RE (%) | CCE (%) | AE (deg) | rMAG (%) |
| --- | --- | --- | --- | --- |
| 5Cb30 | $5.1 \pm 0.6$ | $0.1 \pm 0.0$ | $1.8 \pm 0.0$ | $100.3 \pm 0.1$ |
| | $4.2 \pm 1.6$ | $1.9 \pm 1.5$ | $1.3 \pm 0.2$ | $100.2 \pm 0.4$ |
| 4Cp20 | $19.6 \pm 0.8$ | $1.5 \pm 0.1$ | $6.9 \pm 0.3$ | $109.1 \pm 1.0$ |
| | $11.7 \pm 1.7$ | $11.3 \pm 4.4$ | $3.6 \pm 0.5$ | $101.8 \pm 1.3$ |
| 4Cp25 | $19.0 \pm 0.8$ | $1.4 \pm 0.1$ | $6.8 \pm 0.4$ | $108.8 \pm 1.0$ |
| | $10.9 \pm 1.7$ | $10.1 \pm 4.3$ | $3.6 \pm 0.5$ | $101.5 \pm 1.3$ |
| 4Cp30 | $19.2 \pm 0.8$ | $1.5 \pm 0.1$ | $7.0 \pm 0.3$ | $108.6 \pm 1.1$ |
| | $11.2 \pm 1.6$ | $10.9 \pm 4.5$ | $3.9 \pm 0.5$ | $101.2 \pm 1.4$ |
| 4Cp40 | $20.1 \pm 0.7$ | $1.7 \pm 0.1$ | $7.7 \pm 0.4$ | $108.0 \pm 1.2$ |
| | $12.4 \pm 1.6$ | $13.3 \pm 5.5$ | $4.7 \pm 0.6$ | $100.8 \pm 1.7$ |
| 3S | $27.1 \pm 1.0$ | $4.0 \pm 0.3$ | $12.1 \pm 0.7$ | $97.2 \pm 1.0$ |
| | $22.5 \pm 2.4$ | $23.5 \pm 6.9$ | $6.7 \pm 0.9$ | $87.1 \pm 3.0$ |
| 1S | $28.0 \pm 1.4$ | $4.2 \pm 0.4$ | $13.5 \pm 1.2$ | $95.1 \pm 1.9$ |
| | $23.6 \pm 2.7$ | $23.6 \pm 7.3$ | $6.7 \pm 0.9$ | $84.9 \pm 3.5$ |

## Footnotes

1 Here we use triangle meshes and linear basis functions, but the approach is straightforward to generalize to other element types.

